# Non-equilibrium remodelling of collagen-IV networks *in silico*

**DOI:** 10.1101/2024.10.10.617412

**Authors:** Billie Meadowcroft, Valerio Sorichetti, Eryk Ratajczyk, Fernanda Pérez-Verdugo, Nargess Khalilgharibi, Yanlan Mao, Ivan Palaia, Anđela Šarić

## Abstract

Collagen IV is one of the main components of the basement membrane, a layer of material that lines the majority of tissues in multicellular organisms. Collagen-IV molecules assemble into networks, providing stiffness and elasticity to tissues and informing cell and organ shape, especially during development. In this work, we develop two coarse grained models for collagen-IV molecules that retain biochemical bond specificity and coarse-grain at different length scales. Through molecular dynamics simulations, we test the assembly and mechanics of the resulting networks and measure their response to strain in terms of stress, microscopic alignment, and bond dynamics. Within the basement membrane, collagen-IV networks rearrange by molecule turnover, which affects tissue organisation and can be linked with enzyme activity. Here we explore network rearrangements via bond remodelling — the process of dynamical breaking and remaking of bonds between network molecules. We then investigate the effects of active (enzymatic) bond remodelling. We find that this non-equilibrium remodelling allows a network to keep its integrity under strain, while relaxing fully over a variety of timescales – a dynamic response that is unavailable to networks undergoing equilibrium remodelling.

## I. INTRODUCTION

Mechanical properties of tissues originate from the constituent cells and their interactions, as well as the surrounding medium: the extracellular matrix (ECM). Basement membrane (BM) is a specialised type of ECM that lines epithelial tissues in all multicellular organisms [1, 2].

The BM’s mechanics, mostly due to networks of collagen IV, likely plays a crucial role in shaping cells and tissues during development, homeostasis and ageing [3, 4]. For example, it has recently been shown that the tissue shape in the developing *Drosophila* egg is instructed by BM stiffness: softer BM at the poles of the egg triggers local changes in cell orientations [5]. BM mechanics have also been related to ageing in humans whereby older BMs are stiffer [6], and are associated with an accumulation of collagen IV [7].

It is thought that the ECM, and the BM in particular, behave viscoelastically under applied stress [8–11]. A viscoelastic material deforms elastically on short timescales, without retaining memory of the deformation history. On long timescales, however, it can flow like a liquid [12]. In polymeric materials, viscoelastic behaviour can be caused by entanglements [13–17] or by the breaking and reforming of physical or chemical bonds, which leads to network remodelling. We hereafter use the term *remodelling* to refer to network rearrangement due to the breaking and reforming of bonds. Polymer networks that remodel in this way have been studied by soft matter physicists for decades, due to their mechanical strength, their processability and their ability to self-heal [18– Examples are materials based on hydrogen bonds [23], metal ligands [24] and dynamic covalent bonds [25– Dynamic bond remodelling is also found in many biological components such as microtubule networks [29, 30] and actomyosin gels [31, 32], where cross-linkers bind and unbind from filaments, and even in multi-organism systems such as fire ant rafts [33].

It is challenging to determine the extent of collagen-IV network remodelling in tissues, but collagen-IV turnover within the ECM could be an appropriate proxy [34]. Turnover, measured as the half life of a collagen-IV molecule within the ECM, is known to play a role in development [35], ageing [36] and disease [37– Vastly different values of collagen-IV lifetimes have been reported in the literature, suggesting that the mechanism for degradation and/or replacement of molecules may vary for different systems [34, 35, 41]. For example, finite lifetimes within a network could be the result of breaking and reforming bonds between collagen-IV molecules, with newly formed molecules possibly replacing the old [42]. Turnover has been associated with changes in tissue shape [42] and in most studies it was found to be intimately linked to enzymes such as collagenases [42– Several studies show that collagen-IV cleaving enzymes are up-regulated or down-regulated during development or disease [45– Linking the activity of enzymes and network remodelling dynamics, therefore, may be crucial for understanding the mechanics and viscoelastic properties of ECM components [11, 48] and its subsequent role in the shape and dynamics of tissues [45, 49].

The mechanics and dynamics of collagen-IV networks are largely understudied, as are the microscopic mechanisms of collagen-IV cleaving enzymes. Conversely, the collagen-IV molecule structure and bonding topology have been well established since the 1990s. The collagen-IV units, which bond to make up a network, are called protomers. A collagen-IV protomer is made up of three distinct domains: the NC1 domain at one end, the 7S domain at the other end, and a 390-nm long triple helix connecting them [50, 51] (Fig. 1A). The single protomers bind one another at their ends and assemble into thin sheets [52] (Fig. 1C). In particular, the NC1 ends of two protomers can come together to form an NC1 bond [53, 54] and the 7S ends of up to 4 protomers can form 7S bonds [55–58] (Fig. 1B). Despite genetic similarities with the much more studied collagen I, the structures formed by collagen I and by collagen IV, as well as their binding mechanisms, are considerably different. Collagen-I protomers form long fibrils that stick laterally to each other, assembling into thick, fairly rigid fibres [59–62], while collagen-IV protomers bind through their ends, not forming fibrils or fibres, but rather thin flexible nets. Although lattice models have been proposed to interrogate collagen-IV network mechanics [63, 64], a model incorporating the known molecular details of collagen IV and its bonding is still missing.

**FIG. 1.**
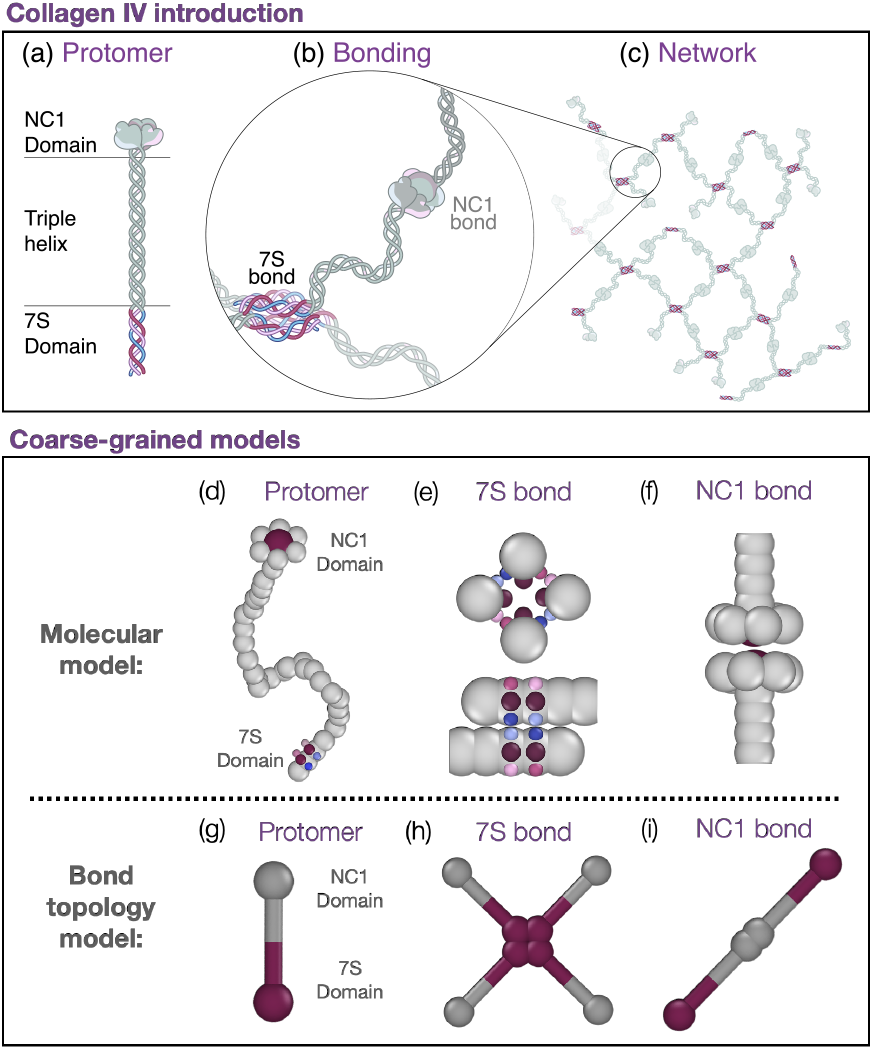
Collagen-IV bonding and models. (**A**) A collagen-IV protomer is made up of a triple helix with an NC1 domain at the C-terminus of the helical chains and a 7S domain at their N-terminus. (**B**) The NC1 end bonds with one other collagen-IV protomer to form an NC1 bond. The 7S end can bond with up to three other collagen-IV protomers to form a 7S tetrameric bond. (**C**) A sheet-like collagen-IV network. (**D**) The molecular model is a beads-spring model where the protomer is made up of 37 beads (with 7 constituting the NC1 domain) connected by springs. The binding regions of the 7S domain are modelled using attractive patches. **(E)** The 7S end forms bonds via 6 interacting patches arranged such that 4 ends can form a stable bond. (**F**) The NC1 end forms bonds via one interacting patch, surrounded by a ring of 6 beads with volume exclusion, to prevent more than two NC1 ends binding. (**G**) In the bond topology model a collagen-IV protomer is represented by two interacting ends connected by a rigid rod with no volume exclusion. (**H**) The 7S end can bond with three other protomers to form the 7S bond. (**I**) The NC1 end can bond with one other protomer to form the NC1 bond.

In this work, we seek to understand how the microscopic properties of the collagen-IV protomer, and most importantly its binding specificity, determine the structure and the mechanical response of the self-assembled networks. We propose two computational models for collagen IV, with different levels of coarse-graining [65]. The first model describes a protomer as a semi-flexible polymer, with ends decorated by specific binding sites (Figs. 1D-F). We use it to study the role of the flexible central chain in network formation. The second model coarse-grains the whole protomer chain to an entropic spring, while retaining bond specificity (Figs. 1G-I). This allows us to reach length scales which are probed in microscopy experiments and to distil how bond kinetics determines macroscopic network properties [3].

To understand the possible role of enzymes in network remodelling, we then compare two different routes for bond rupture and formation, using the coarser bond topology model. One route, termed the equilibrium protocol, locally obeys detailed balance [66], while the other route, termed enzymatic protocol, does not. We find that active (enzymatic) remodelling brings the network to relaxed states which are forbidden under equilibrium remodelling. Our results confirm that active processes strongly affect the structure and mechanics of self-assembled materials [67–71].

## II. MODELS AND SIMULATION METHODS

Here we present the two particle-based models, each simulated using molecular dynamics.

The “molecular model” represents a protomer (including the triple helix and the binding domains) as a semiflexible chain made of spherical beads connected by harmonic springs (Fig. 1D). Short-range attractive patches ensure the 7S end of the protomer bonds with two antiparallel protomers on their 7S ends, which in turn bond with a fourth parallel protomer on its 7S end, as proposed in the literature [55, 72] (Fig. 1B). This results in the formation of dimers, trimers, or at most tetramers, as depicted in Fig. 1E [58]. The attractive site at the other end, protected by six inert beads to ensure one-to-one binding, represents the bulkier NC1 domain and bonds with the NC1 end of another protomer, forming the dimer depicted in Fig. 1F [53, 54]. Volume exclusion ensures no chain crossing can take place, so that entanglement between chains is correctly captured.

The molecular model, as per its name, has the advantage of providing a more direct connection to the molecular details of the system. However, despite its simplicity, simulating a system size that can be directly compared to conventional microscopy data is still not feasible due to prohibitively large computational times.

To simulate networks which are closer to the typical experimental sample sizes and time scales [3], we developed the “bond topology model” which represents each molecule using two beads only. This is reminiscent of other recent efforts at reducing the complexity of elastomer network models by mapping full polymer systems to simplified mesoscale models [22]. Few parameters define the length and energy scales of the protomer and its bonding within this model, and each is chosen in such a way to coarse-grain a key aspect of the molecular model. The parameter choice is discussed in detail in sec. I of the SI.

In the bond topology model, a protomer is represented as a two-bead rod, as shown in Fig. 1G. The 7S bead can bond with beads of the same kind from up to three other protomers (Fig. 1H), while the NC1 bead can bond with the NC1 bead of only one other protomer (Fig. 1I). Rods are treated as rigid bodies and the flexibility of the chain is effectively encoded in the harmonic potentials that make up the 7S and NC1 bonds. The bonding distance is imposed by a two-body harmonic potential, whereas a three-body harmonic potential controls the angular flexibility of the bonds between two protomers. A collagen network is therefore defined only through its bond topology and through the distance that protomers impose between binding sites.

For both models, parameters are chosen such that the majority of the protomers spontaneously assemble into a percolating network. The network structure is, however, not fixed: bonds are dynamic and can break and reform due to thermal fluctuations. We first consider passive remodelling (sections A and B). In the molecular model, bonds are formed and broken through attractive interactions, according to the equilibrium canonical distribution using molecular dynamics. The strength of the attraction determines the bond remodelling dynamics. In the bond topology model, molecular dynamics is coupled with a Metropolis algorithm [73] that explicitly breaks and forms bonds locally respecting detailed balance, thus changing connectivity but still sampling the equilibrium distribution (see Fig. 4A). Then (in section C), we introduce active bond remodelling for the bond topology model. For this purpose we modify the algorithm in breach of detailed balance, so that bonds break and form either randomly or with a distance-dependent rule, as further explained in section III C (see Fig. 4B).

Particle motion is the result of overdamped Langevin dynamics [74], so that the solvent is treated implicitly and hydrodynamics is neglected. Simulations are performed in LAMMPS [75] and visualised using OVITO [76]. The REACTER package [77] was used for the bond topology model. Further details on the models, simulation setup and parameters are outlined in sec. I of the SI.

## III. RESULTS

### A. Self-assembly

We first set out to explore how the collagen-IV protomers self-assemble into networks, in absence of external strain. The simulations begin with the protomers in a random configuration, from which they freely diffuse and bond/unbond with other protomers until the system reaches a steady state, as shown in the snapshots, Figs. 2A and D and SI Videos 1 and 2.

**FIG. 2.**
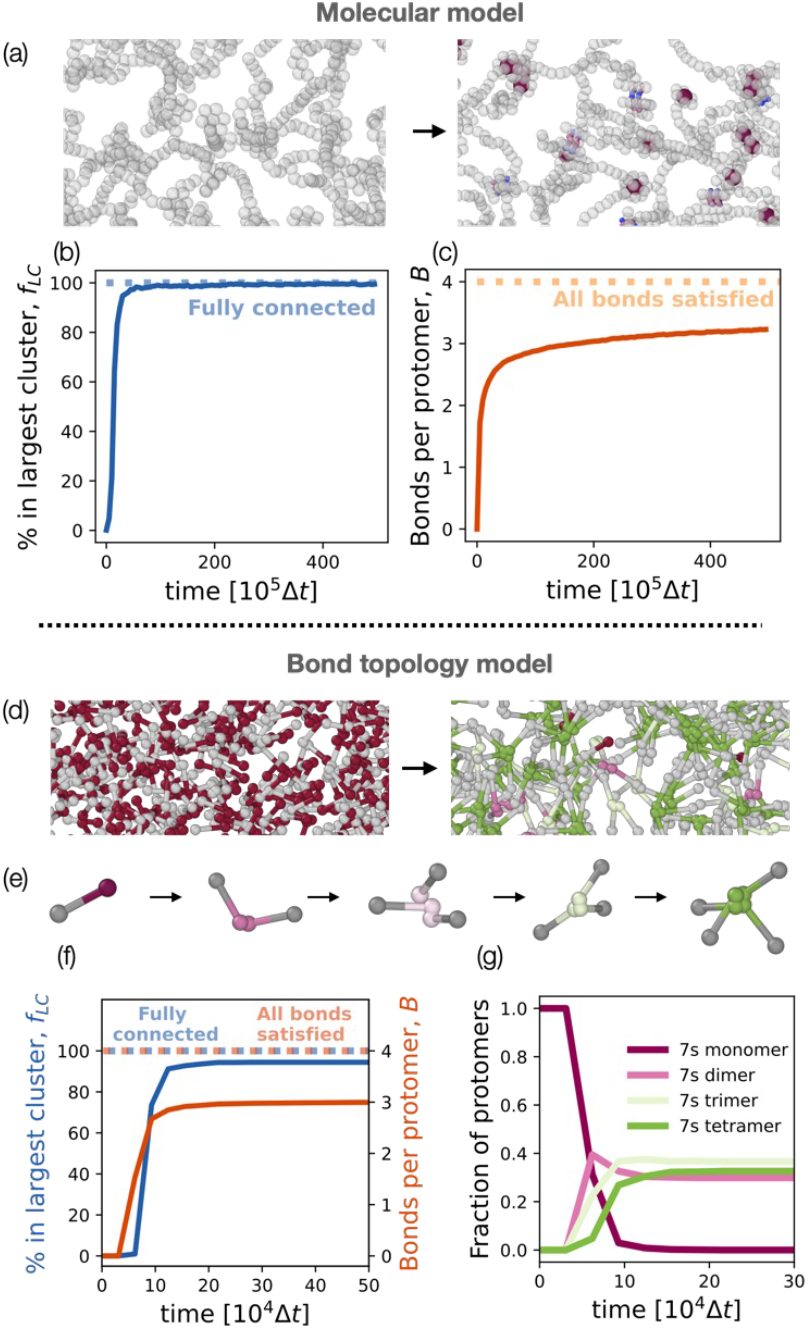
Network self-assembly. The molecular model and the bond topology model both self-assemble into percolating networks. (**A**) Snapshots of the molecular model before (left) and after (right) the network self-assembles. The molecule ends are shown with colour if they are bonded (coloured as in Fig. 1D-F). (**B**) The fraction of molecules in the largest connected network (*f*_*LC*_) over time. The network percolates after a time *≈* 5 *×* 10^6^Δ*t*. (**C**) The number of bonds per protomer in time, which saturates to *≈* 3.2. If all bonds were satisfied the number of bonds per protomer would be 4 (1 from the NC1 domain and 3 from the 7S domain). (**D**) Snapshots of the bond topology model before (left) and after (right) selfassembly. (**E**) The pathway for the bonding of 7S ends, which are coloured according to the degree of bonding as shown in the legend G. (**F**) The fraction of molecules in the largest cluster and the bonds per protomer in time. The network percolates after *≈* 1.0 *×* 10^5^Δ*t* and the number of bonds per protomer saturates to 3. (**G**) The time evolution of the formation of 7S bonds by number of bonding partners. The 7S dimer bonds are the first to form but over long times the 7S tetramer and trimer bonds dominate.

We analyse the connectivity of these networks by calculating the fraction of protomers in the largest connected cluster, *f*_*LC*_, which gives a measure of network connectivity. In both models the binding energy of the protomer-protomer bonds are varied to identify the percolation threshold. We thereafter choose bond energies to be above this threshold such that we only investigate percolating networks (see Fig. S9). Figs. 2B and F show *f*_*LC*_ as a function of time for networks that become fully connected. Note that the simulation time step Δ*t* corresponds to a larger physical time in the case of the more coarse-grained, lower density, bond-topology model (see sec. IV of the SI).

Both models form networks that on average form a total of *B* = 3 − 3.2 bonds per protomer, as shown in Figs. 2C and F. This is significantly below the value of *B* = 4 which would indicate that all bonds are satisfied and corresponds to the system’s ground state. Experimental studies of collagen-IV assembly have indicated a value *B* ≈ 3.2 bonds per protomer from *in vitro* low-concentration essays [58] and *B* ≈ 3 from eye tissue imaging [78]. However, the connectivity of collagen-IV networks within the basement membrane is unknown. In our models, the value *B* ≈ 3 − 3.2 bonds per protomer is maintained when bond strength is increased (Figs. S9C and D), suggesting that the system cannot explore all bond configurations due to kinetic trapping in local free energy minima.

In the bond topology model, we additionally monitor how the 7S tetramer bonds form. Fig. 2E shows typical snapshots of the 7S bonding pathway. In Fig. 2G we see ≈70% of tetramers and trimers make up the percolating network, while ≈30% of the 7S bonds are still in the dimer configuration. Overall, these result validate the two models, which are in line with each other and corroborate the available experimental data.

### B. Mechanics

To investigate the mechanics of the self-assembled networks, we apply a step strain and monitor the response of the networks to this strain. The strain is applied over a short time interval and over one direction, similar to those applied in experiments probing mechanics of tissues in fly wing discs. For example, manual stretchers have been used to strain the wing disc by up to ∼ 100%, while the cells can be imaged as they respond to the perturbation [79]. The strain protocol is represented in Figs. 3A-B (see also sec. II of the SI for details). The snapshots in Figs. 3C-D show the networks before, immediately after and some time after the strain is applied, also shown in SI Videos 3 and 4. The macroscopic mechanical response of the network is studied by measuring stress in the direction of strain. The microscopic response is examined by measuring the individual protomer alignment with the direction of strain and by characterizing the bond kinetics.

**FIG. 3.**
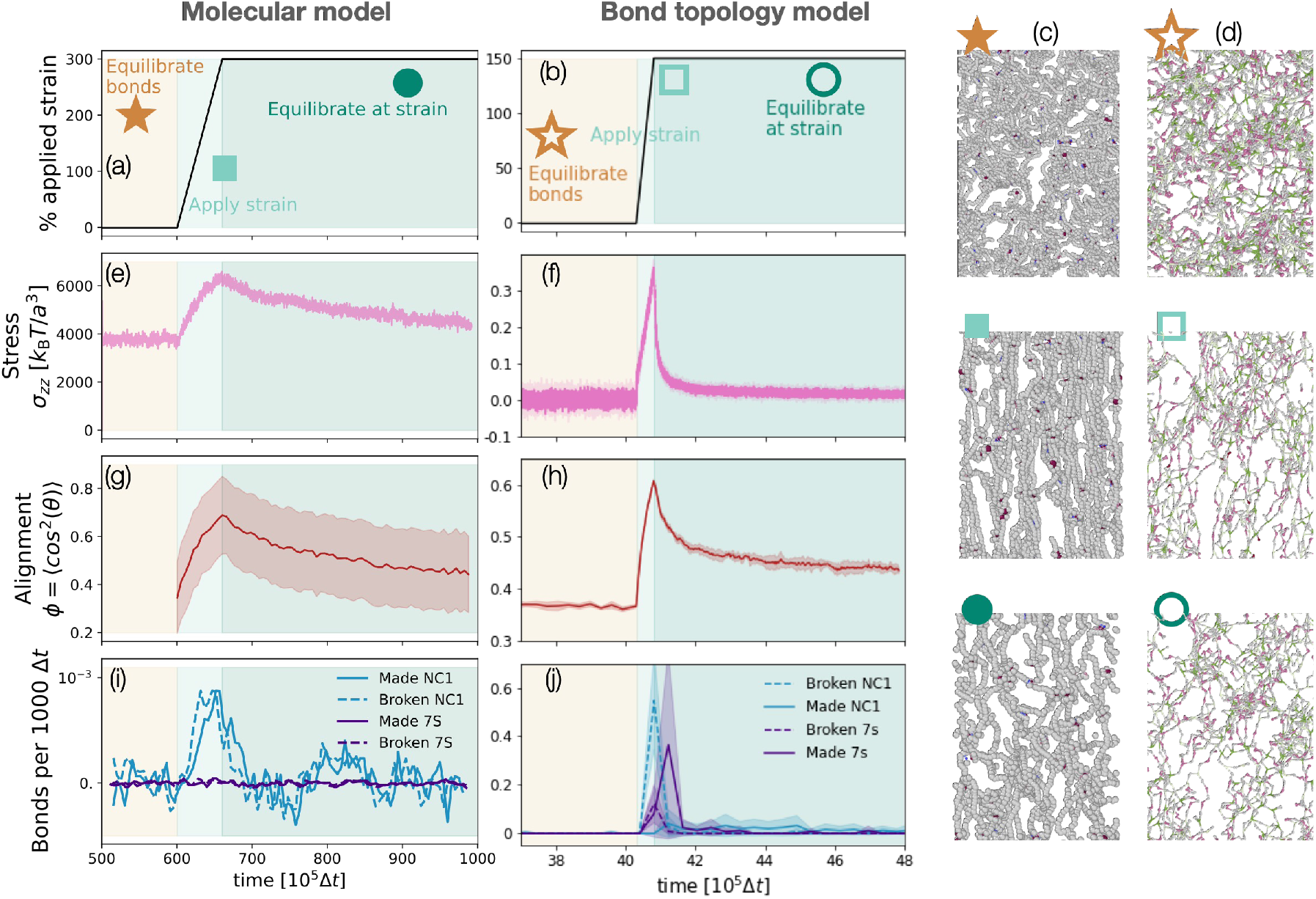
Mechanical response to strain. (**A-B**) The networks are subject to strain after equilibration. (**A**) The molecular model is subject to a strain of 300% in the *z*-direction, which is applied at constant volume. (**B**) The bond topology model is subject to a strain of 150% in the *z*-direction where the box size is kept constant in the *x*- and *y*-directions. (**C**) Snapshots of the molecular model and (**D**) of the bond topology model before strain is applied (⋆), immediately after strain is applied (▪) and after network relaxation (). (**E**-**F**) The stress in the direction of strain over time in the molecular model and bond topology model. The stress reaches a peak at the end of the strain application and then relaxes while the strain is kept constant. (**G**-**H**) Protomer alignment with respect to the direction of strain, averaged over all protomers. (**I**-**J**) The number of bonds broken and made per 1000 simulation time steps. In both models the ratio of bonds broken to bonds made increases when the strain is applied. (**I**) The bonds broken in the molecular model increase sharply when strain is applied which is followed by an increase in bonds made as the ends from previously broken bonds form new ones. (**J**) The number of bonds broken in the bond topology model also increase at peak stress, followed by an increase in number of bonds made.

**FIG. 4.**
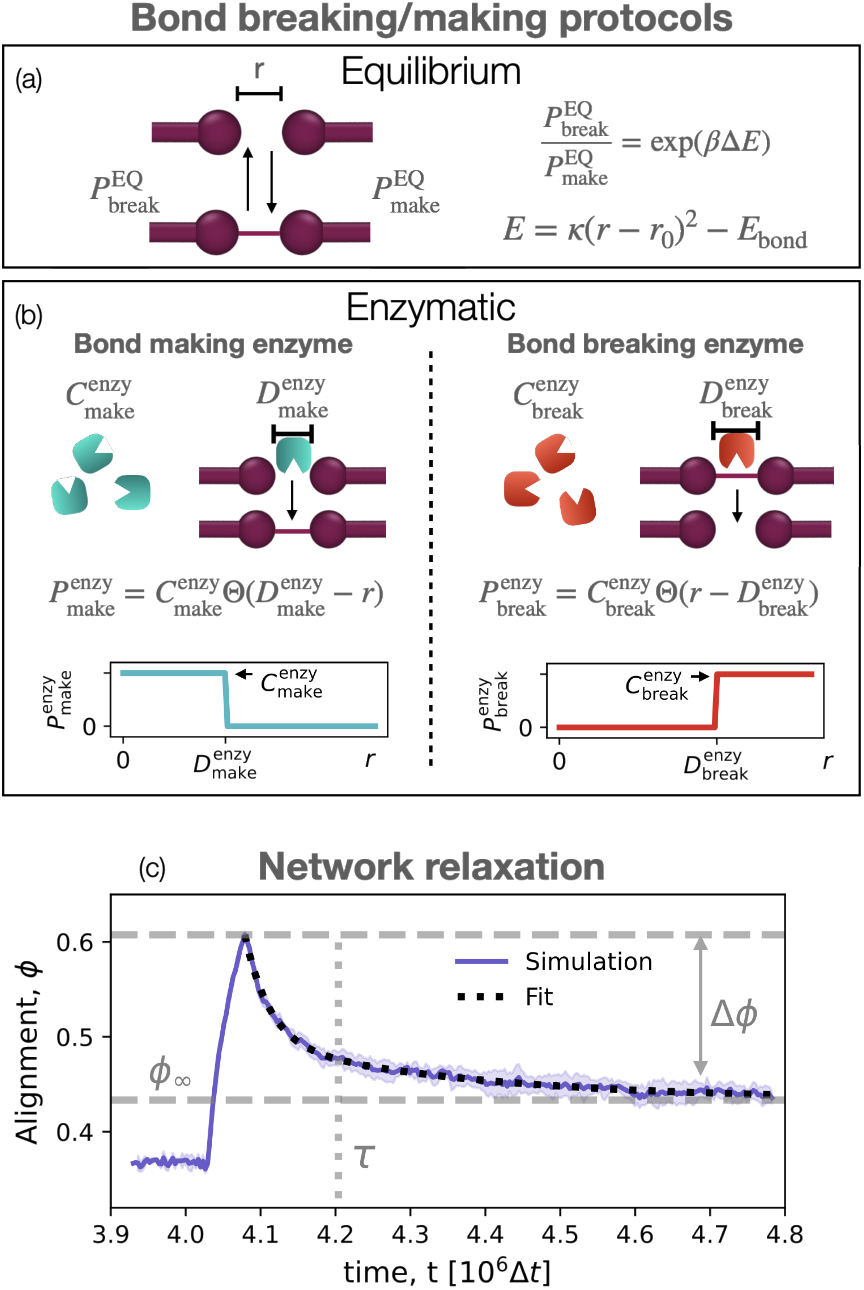
Bond remodelling protocols and network relaxation. (**A**) The bond breaking and making protocols in the equilibrium regime. The bonds break and make with probabilities that satisfy detailed balance, Eq. (1) (**B**) The bond breaking and making protocols in the non-equilibrium/enzymatic regime. The bonds make or break with probabilities which are proportional to an effective bond-making or -breaking enzyme concentration, 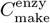 or 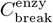. The fictive enzyme is only active when two ends are within the enzyme activation range, defined by distances 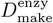 and 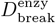. **(C)** Network alignment relaxation in response to strain, with the fit as described in Eq. 4.

The stress response *σ*_*zz*_, with z being the direction of strain, is shown in Figs. 3E-F. In both models, the stress peaks once the final strain is reached and then decreases over time as the strain is kept constant. This underlies the viscoelastic nature of the material, with an elastic response at small time scales (linear increase of stress upon linear increase of strain) followed by a viscous relaxation at large time scales (stress relaxation at constant strain). The stress relaxation of the material is well described by a two-mode generalised maxwell model [12], *i*.*e. σ*_*zz*_(*t*) = Δ*σ*_*zz*_(*f* × exp(−*t/τ*_1_)+(1 − *f*) × exp(−*t/τ*_2_)), where Δ*σ*_*zz*_ is the increase of stress upon stretch, *τ*_1_ is the fast relaxation mostly due to protomer rearrangement in space and *τ*_2_ is the slow relaxation due to the bond remodelling. We further characterise the Young modulus and the critical strain, the onset of strain-stiffening [80], of the bond topology model networks in sec. III of the SI. A small-strain analysis results in a Young modulus that is small (≈ 0.03 Pa) compared to measurements of 3D networks of other ECM components [8]. This possibly reflects the thin semi-2D nature of the networks simulated here, and suggests that the non-collagen-IV components of the BM might play a key role at small strain or that the *in-vivo* collagen-IV matrix might be pre-stressed during development.

To gain insight in how viscoelasticity emerges from the microscopic rearrangement of protomers, we look at single protomer alignment, which we define by cos^2^ *θ*, where *θ* is the angle between the protomer end-to-end axis and the direction of strain. We measure the average alignment over all protomers, *ϕ* = ⟨cos^2^ *θ*⟩, in Figs. 3G-H. Completely random alignment corresponds to *ϕ* = 1*/*3. We see that the macroscopic stress response is reflected in the microscopic alignment response: the alignment also peaks when the final strain is reached and slowly decays as the strain is kept constant. From the qualitative agreement between the molecular model and the bond topology model we deduce that entanglement between molecules, which could slow down dynamics and stress relaxation by orders of magnitude [13, 15, 17, 81], is negligible in the molecular model, at least for these densities. This justifies the choice to neglect excluded volume effects in the bond topology model.

As both models have non-permanent bonds, we examined whether there is a change in the bonding dynamics in response to the applied strain. Figs. 3I-J show measurements of the bonds broken and made per time step for each model. In both cases, when strain is applied, temporarily more bonds are broken than made. Balance is quickly restored, as new, unstressed, bonds are formed. Taken together, the networks behave elastically on short time scale in response to external strain, generating stress. This stress induces sudden but persisting align-ment of protomers in the direction of strain and is successively relieved, on longer time scales, by bond breaking and remaking. These results connect the macroscopic viscoelastic response of the networks to the microscopic network arrangement and bond remodelling dynamics. The stress relaxation mechanism measured here is analogous to that observed in other polymer networks with reversible bonds [21, 22, 28]. In particular, the correlation between stress and molecular alignment relaxation was also reported in simulations of vitrimers [28].

As the resolution of microscopes improves, the alignment of molecules within tissues is likely to become more easily measurable [82, 83]. We therefore expect this link between molecule alignment, stress and bond dynamics to become a powerful tool to probe microscopic properties of tissues.

### C. Equilibrium Vs Enzymatic remodelling

In developing organisms the collagen-IV networks in the BM are subject to stress during growth, but the BM is not isolated. Instead, it is always in contact with cells, which actively secrete proteins and enzymes. Some of these enzymes are known to be involved in collagen-IV remodelling and degradation [43, 44, 84]. Here, we explore how the mechanical relaxation and integrity of networks subject to active, enzymatic-like bond remodelling differs from those undergoing equilibrium bond remodelling. In doing so, we restrict ourselves to the bond topology model, because the larger accessible system sizes and time scales make the results more readily comparable with experimental observations of the basement membrane collected in light microscopy imaging, and allow us to reduce data noise.

The equilibrium bond remodelling protocol satisfies detailed balance, meaning that the ratio between the probabilities of making and breaking a bond obey

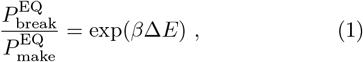

where *β* = 1*/*(*k*_B_*T*), *k*_B_ is the Boltzmann constant, *T* is temperature, and Δ*E* is the bond energy (Fig. 4A). This relation does not need to hold true for the enzymatic bond remodelling regime where probabilities may depend on other factors, like effective enzyme concentrations and local conformation of the binding sites. For example, mechano-sensitive enzymes have been suggested to regulate many other biological filaments and networks such as ESCRT [85] and actin [86]. Additionally, enzymes from the family of matrix metalloproteinases, known to degrade different types of collagen, have been found to act in a strain-dependent manner [87–89]. The non-equilibrium protocol we choose takes inspiration from these mechano-sensitive enzymes, which may have different affinities for stretched and relaxed target domains. We model enzymatic remodelling assuming the following probabilities for bond making and breaking

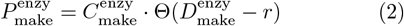

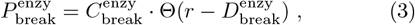

where 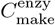 and 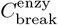 are effective parameters representing the concentration of bond-making and -breaking enzymes, Θ is the Heaviside step function, and *r* is the bond length – a probe for the binding site strain. Enzymes therefore create bonds when two ends from different protomers come closer than the activation distance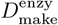 and break bonds when two ends are stretched further apart than the effective activation distance 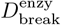 (Fig. 4B).

We quantify the relaxation dynamics by measuring the average molecule alignment in the direction of strain, *ϕ* = ⟨cos^2^ *θ*⟩, a robust and relatively noise-free measurement (see Fig. 3H). Similar to the stress relaxation, the alignment response follows a double exponential decay, shown as the fitting curve in Fig. 4C:

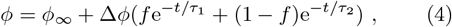

where relaxation times *τ*_1_ and *τ*_2_, the amplitude ratio, *f* and the long-time residual alignment *ϕ*_∞_ are fitting parameters. The amount of relaxation Δ*ϕ* is measured from the alignment curves. We define the overall relaxation time as the amplitude-weighted relaxation time *τ* = *f* × *τ*_1_ + (1 − *f*) × *τ*_2_. Reducing the relaxation response to these parameters allows us to summarise the effects on relaxation of the bond remodelling parameters and protocol.

When bonds are remodelled following the equilibrium protocol, we find a strong correlation between the amount of network relaxation, Δ*ϕ*, and the percentage of protomers *not* in the largest cluster at long times, 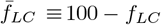, which is a probe for network fragmentation.We will refer to this quantity as “fragmentation” for simplicity. In particular, the amount of relaxation scaled by the strain 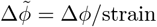, which we hereafter call simply relaxation amount, collapses to a single line when plotted against fragmentation (Fig. 5A). The networks are less fragmented if the chemical bond energy, *E*_bond_, is larger (shown in different colours in Fig. 5A), and this results in a higher initial alignment of the networks and therefore higher 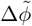. This scaling holds across different applied strains, which are represented by different symbol shapes in Fig. 5A. Additionally, the amount of network relaxation is also correlated with the speed of relaxation: the networks with higher bond energies, *E*_bond_, have slower bond remodelling dynamics and therefore take longer to relax. This is shown in Fig. 5B and is again independent of the amount of strain.

**FIG. 5.**
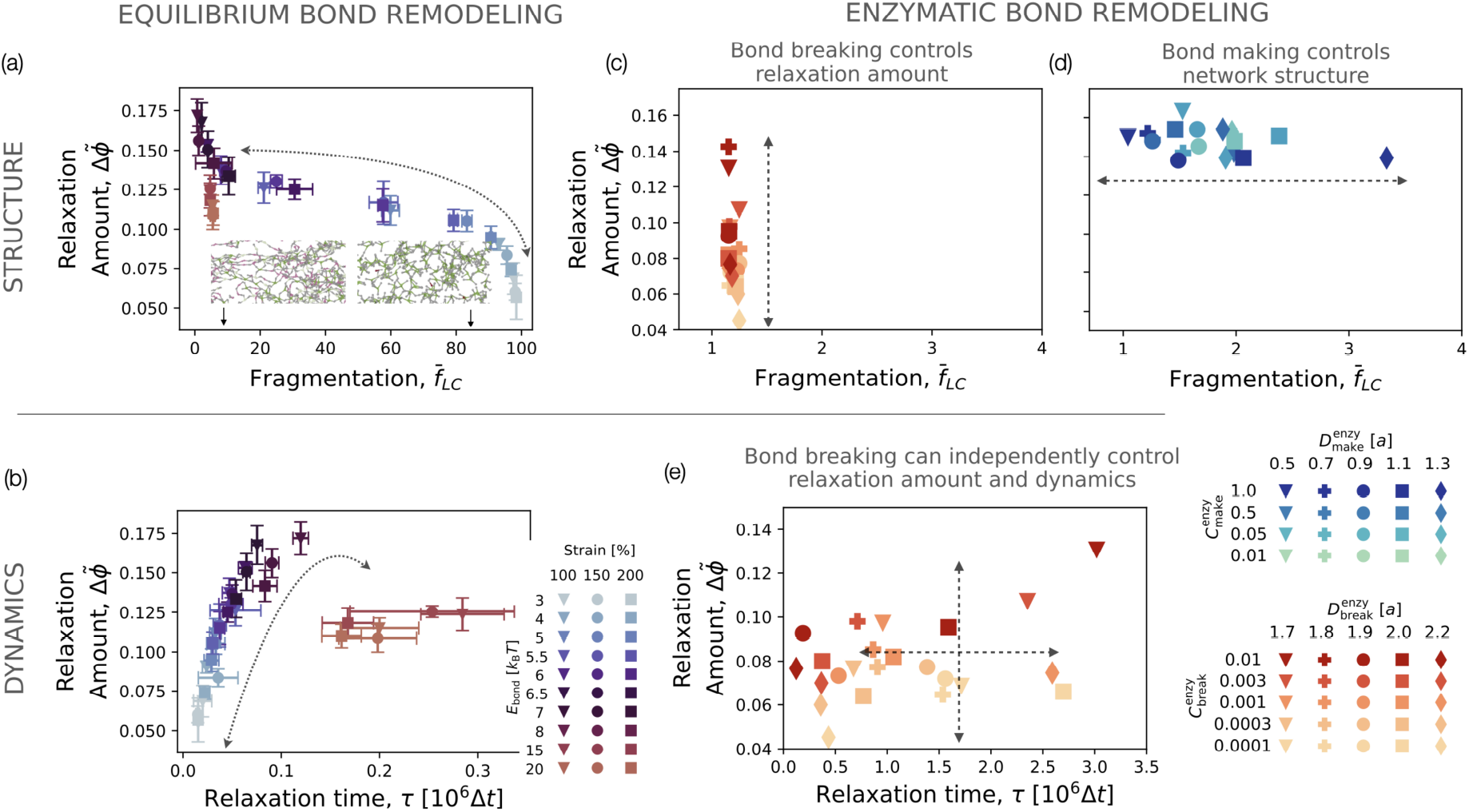
Relaxation dynamics for equilibrium and enzymatic bond remodelling. (**A** and **B**) Relaxation dynamics in the equilibrium regime. (**A**) The amount of network relaxation scaled by the strain, 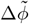, is plotted against the percentage of the network not in the largest connected cluster, 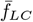, for different amounts of strain and chemical bond energies, *E*_bond_. When 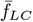 is 100%, the network is completely fragmented, see bottom snapshots for networks with low (left) and high (right) fragmentation. In the equilibrium bond remodelling regime, the amount a network can relax, 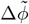, is larger for lower 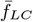 and visa versa. The error bar is the standard deviation calculated from 5 random seeds. (**B**) In the equilibrium bond remodelling regime, 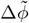 is proportional to the relaxation timescale, *τ*, for small bond energies, after which the relaxation timescale does not change. (**C** - **E**) Relaxation dynamics in the enzymatic regime, for the influence of bond breaking, (**C**) and (**E**) and bond making, (**D**). (**C**) The amount of relaxation scaled by strain, 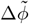, is plotted against the network fragmentation, 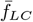, for varying bond breaking parameters where different 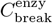 are represented by different colours and different 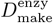 are represented by different symbols. The trend shows that when the bond making parameters are kept fixed, the bond breaking parameters (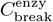 and 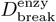) do not influence the fragmentation of the network. (**D**) In the enzymatic regime, 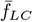 can be shifted to low values by changing the bond making parameters, 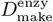 and 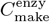, which does not affect the amount of relaxation, 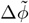. (**E**) The amount of network relaxation, 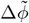, is plotted against relaxation timescale, *τ*, for the bond breaking parameters as in (**C**). This plot shows that 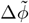 and *τ* can be tuned independently using enzymatic remodelling. Roughly, different 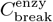 can tune the amount of relaxation 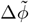 and different 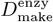 can tune the speed of relaxation, *τ*. For (**C**) and (**E**) the strain is 150%, 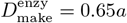 and 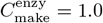. For (**D**) the strain is 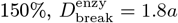, and 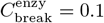.

We note that equilibrium networks fall into two distinct regimes. Those with *E*_bond_ ≲ 7*k*_B_*T* undergo continuous thermally driven bond rearrangement, while those with *E*_bond_ ≳ 7*k*_B_*T* are kinetically trapped and rearrange only under applied stretch. If collagen-IV networks remodel via equilibrium mechanisms, they likely fall into the latter regime, as their covalent and disulphide bonds exceed 7*k*_B_*T* [90]. This is in line with other observations of collagen-IV networks *in vitro*, which have ∼3.2 bonds per protomer [58] — a value we observe in the high bond energy limit. In contrast, low bond energy networks show a higher number of bonds per protomer (Fig. S9D).

The enzymatic regime deviates from the equilibrium regime in two significant ways. Firstly, while in equilibrium the bond dynamics is constrained by the bond energy through detailed balance as per Eq. (1), out of equilibrium bond making and bond breaking are fully independent and regulated by distinct quantities. We observe that relaxation dynamics is essentially determined by bond breaking enzymes, whereas bond making enzymes governs network connectivity. This means that within enzymatic remodelling the correlation between connectivity and the amount the network relaxes, seen in Fig. 5A for the equilibrium regime, does not have to hold.

This is evident in Fig. 5D, where by increasing the bond making activation distance, 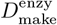 (symbols) or the concentration of bond making enzymes, 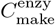 (colours), the system deviates from the connectivity prescribed by equilibrium. Additionally, varying the bond breaking parameters determines the amount of relaxation, 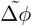, independently from connectivity, see Fig. 5C. This means that enzymatic bond remodelling allows stress to relax without compromising the structural integrity of the network. Varying the concentration of breaking and making enzymes, one can cover more of the phase space represented in Figs. 5C and D without fragmenting the network, whereas at equilibrium the available phase space is constrained (Fig. 5A).

The second way in which the enzymatic regime deviates from the equilibrium regime is more subtle. In the equilibrium regime, one parameter controls the frequency of bond breaking: the bond energy. In the enzymatic regime, there are two parameters contributing to this frequency: the bond breaking probability, 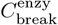 and the bond breaking activation distance, 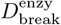. The overall frequency of bond breaking is proportional to 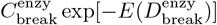, where the first term is the probability of a bond within activation distance to break and the second term is the probability of being within activation distance (modulo a constant). Therefore, the same overall bond breaking frequency can be obtained by different combinations of the bond breaking parameters 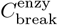 and 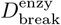, meaning that the system can be more or less sensitive to how stretched bonds are.

When an enzyme is more sensitive to bond length and only breaks stretched bonds, networks relax faster: the breaking of stretched bonds, as opposed to that of unstretched bonds, drives stress-relaxation. This is clear from Fig. 5E where the different colours represent the enzyme concentration, and the different symbols represent the bond breaking activation distance. In this graph, the colours dictate the amount of relaxation while the symbols separately determine the timescale of relaxation.

Once more, varying the breaking parameters allows the tiling of the whole phase space in Fig. 5E, unlocking relaxation pathways that are unachievable in equilibrium (Fig. 5B). In summary, in the enzymatic regime the relaxation timescales can be tuned independently of the amount of relaxation, while in the equilibrium case these are intrinsically coupled.

It is possible to implement the bond remodelling in the enzymatic regime such that it is non-mechano-sensitive. This is the extreme case where the bond breaking activation distance 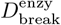 becomes zero and any bond may be broken. The enzyme concentration in this case is the only parameter dictating the bond breaking dynamics. The results for non-mechano-sensitive bond remodelling are shown in Fig. S10, showing that these networks can also relax with a range of timescales and amounts while keeping fully connected.

## IV. DISCUSSION

We presented the first computational framework to investigate how the mechanics and dynamics of the basement membrane may be influenced by collagen-IV bond remodelling and the activity of enzymes. By developing two models of collagen IV with two different levels of coarse-graining, we investigate the connectivity of percolating collagen-IV networks and link this to experimental observations. We then applied a step strain to the networks to measure the stress response. We find that in the regime of high strain the relaxation dynamics in both models are dominated by bond remodelling, meaning the networks relieve stress by breaking and reforming bonds. Within this regime, we systematically explore different protocols for bond remodelling using the more coarsegrained model. We compare equilibrium and enzymatic bond remodelling protocols, where enzymatic protocols are not constrained to satisfy equilibrium thermodynamics. A key finding is that in equilibrium, there is an inter-dependence between the amount of stress relaxation the network can undergo, its structural integrity and the typical relaxation time. In other words, these different physical quantities cannot be tuned independently. Enzymatic bond remodelling, however, is not restricted in such a way, so that relaxation pathways are available that are inaccessible in equilibrium.

This is a powerful result showing that materials forming and developing while consuming energy have different and more diverse mechanical properties than their passive counterparts. Previous studies on various systems have demonstrated that structures assembled out of equilibrium show a signature of broken detailed balance through non-zero currents in phase space, large-scale fluctuations, or unusual mechanical responses, even once activity has stopped [68, 70, 71, 91]. In general there are many possible ways of identifying non-equilibrium signatures. Here we show that an under explored approach is to analyse the relations between network fragmentation and relaxation dynamics. This new metric for non-equilibrium is particularly promising, as it can in principle be checked for in experiment; it relies only on measuring how alignment varies in time under strain, and might allow us to infer microscopic remodelling mechanisms from macroscopic observations.

The concentrations and mechano-sensitive properties of bond-cleaving enzymes emerge as free parameters and are capable, in principle, of tuning the structure and mechanics of the network. This may be crucial in biology, especially in developing organisms where large shape changes must occur in a controlled manner [92]. For example, the collagen-IV network in the basement membrane of a developing tissue provides mechanical stiffness on short timescales, but must relax on the timescale of tissue growth [41]. Using enzymes, an organism could achieve precise timescales of relaxation and amount of relaxation, while keeping network integrity. Additionally, during development the basement membrane properties may change in time by simply changing enzyme activity, rather than needing to change the chemical nature of the network itself. Without enzymatic activity, the material is restricted to mechanical properties which are solely governed by the chemistry and connectivity of the underlying network. We envisage that a future direction of this work will be to further understand the changes in basement membrane mechanics due to ageing [36, 93, 94], disease [95] or drugs [96], which all could be due to changes in enzyme activity and collagen-IV turnover.

It should be noted that there are numerous ways in which active systems may deviate from equilibrium bond remodelling kinetics, and we chose here just one, where enzymes actively break and make bonds depending on bond distance or else randomly. Other examples might include the action of bond swapping, whereby the number of bonds in the network stays strictly constant but energy is consumed to swap bonds between neighbours, similar to vitrimer networks [27]. Another mechanism is present in actively cross-linked bio-networks like actomyosin, where structural changes may be caused by bond-sliding using the action of walking motors [31].

We here chose a simple realisation of bond remodelling,where both NC1 bonds and 7S bonds can be broken and made, and the triple helix chain is never broken. This assumption is likely valid for equilibrium bond remodelling because the intra-protomer bonds of the triple helix are more stable than the inter-protomer bonds [97]. However enzymes like collagenase could degrade collagen-IV networks in other ways and may well degrade the triple helix [98] which would be less likely to rebind into a network. We note that if enzymes degrade the protomers’ binding domain, such that the second neighbours in the network are free to rebind [42], the model proposed here remains a good approximation. We break bonds one at a time, though the results would not be significantly different if we were to break the two protomer bonds simultaneously.

The methods and models introduced here should be considered as a minimal framework to explore and contrast the active and passive remodelling of collagen-IV networks. Systematically exploring other non-equilibrium bond remodelling regimes and their effects on the mechanics of networks is a compelling future direction. This development would be in tandem with ever improving experimental techniques in mechanobiology which are holding great promise to elucidate the non-equilibrium mechanism behind bond remodelling of collagen IV *in vivo*.

## V. ACKNOWLEDGEMENTS

This work has received funding from the European Research Council (ERC) under the European Union’s Horizon 2020 research and innovation programme (grant agreement No. 802960; B.M., V.S., I.P., A.Š.). I.P. also acknowledges funding from the European Union’s Horizon 2020 research and innovation programme under the Marie Skłodowska-Curie grant agreement No. 101034413. F.P.V. acknowledges support from the NOMIS foundation. We thank the National Centre for the Replacement, Refinement and Reduction of Animals in Research (NC3Rs) for funding this work (NC/T002425/1 to N.K.). N.K. was also supported by a Leverhulme Trust project grant (RPG-2020-068) and an MRC Fellowship (MR/W027437/1) awarded to Y.M. Y.M. was funded by MRC award MR/W027437/1, a Lister Institute Research Prize and EMBO Young Investigator Programme.

