## Supplementary Information for "Non-equilibrium remodelling of collagen-IV networks *in silico*"

### Supplementary Information
### Nonequilibrium Remodeling of Collagen-IV Networks *in Silico*

Billie Meadowcroft,[1,2,3,4] Valerio Sorichetti,[1] Eryk Ratajczyk,[4,5,6] Fernanda Pérez-Verdugo,[1] Nargess Khalilgharibi,[2,3] Yanlan Mao,[2,3] Ivan Palaia,[1,7] and Anđela Šarić[1]

[1]*Institute of Science and Technology Austria, Klosterneuburg, Austria*
[2]*UCL Laboratory for Molecular Cell Biology, University College London, Gower Street, London WC1E 6BT, United Kingdom*
[3]*Institute for the Physics of Living Systems, University College London, Gower Street, London WC1E 6BT, United Kingdom*
[4]*Department of Physics and Astronomy, University College London, London WC1E 6BT, United Kingdom*
[5]*Rudolf Peierls Centre for Theoretical Physics, University of Oxford, Parks Road, Oxford OX1 3PU, United Kingdom*
[6]*Kavli Institute for Nanoscience Discovery, University of Oxford, South Parks Road, Oxford OX1 3QU, United Kingdom*
[7]*Department of Physics, King's College London, WC2R 2LS, United Kingdom*

(Dated: August 4, 2025)

#### I. MODEL DETAILS

##### A. Molecular model

**Model** In the molecular model (see Fig. S1 and Fig. 1D in the main text), a protomer chain, representing the triple helix (including the 7S domain), is made up of 30 beads. The beads are connected by harmonic bonds with energy $E_{\text{len}} = \kappa_{\text{len}}(r - r_0)^2$ where $r$ is the distance between two beads, $r_0 = 0.5a$ is their equilibrium distance where $a$ is the simulation length unit and $\kappa_{\text{len}} = 50k_{\text{B}}Ta^{-2}$ is the bond spring constant, chosen so that bonds are stiff. Including the central bead of the NC1 domain (see below), the contour length of the protomer is $\mathcal{L} = 31 \times 0.5a = 15.5a$. Since experimentally $\mathcal{L} \approx 400$ nm [1], our simulation length unit $a$ maps to approximately 26 nm.

The bending rigidity of the chain is set by the angular potential $E_{\text{ang}} = \kappa_{\text{ang}}(\beta - \beta_0)^2$, where $\beta$ is the angle between three consecutive chain beads in a protomer and $\beta_0 = 180°$. The chain section close to the NC1 domain is $\approx 10$ times more rigid than the rest of the chain which reflects experimental observations [1] (see Fig. S1). However, implementing a uniform persistence length results in qualitatively the same mechanics as the heterogeneous protomer, as shown in Fig. S2. The chosen angular potentials gives a persistence length of the chain $L_p = 1.8a$, approximately 11.6% of the contour length of the protomer. This is consistent with the experimentally measured value of $L_p$ as 10% of the contour length [1]. Section V details the calculation of the persistence length.

All beads in the model interact via a truncated and shifted Lennard-Jones potential:

$$E^i_{\text{LJ}}(r) = 4\epsilon^i_{\text{LJ}} \left[ \left(\frac{a^i}{r}\right)^{12} - \left(\frac{a^i}{r}\right)^6 \right] + \alpha \quad \text{for } r < r^i_c, \quad \text{(S1)}$$

$\alpha$ takes its value to ensure $E^i_{\text{LJ}}(r^i_c) = 0$ and $i$ denotes the type of bead.

In addition to the harmonic springs connecting consecutive chain beads of the same protomer, the interaction between any bead pair excluding the attractive patches (described in the next paragraphs) is purely repulsive ($r^{\text{chain}}_c = 2^{1/6}a$), with $\epsilon^{\text{chain}}_{\text{LJ}} = 2k_{\text{B}}T$. This is chosen to be large enough to ensure that chains to not intersect, yet ensures numerical stability with a reasonable time step. To increase the simulation efficiency, the repulsive interaction between first, second and third bonded neighbors in the protomer chain is turned off, which is justified by the fact that they never come into close contact.

The 7S domain is made up of 6 attractive patches on the last three beads of the chain (Fig. S1 and Fig. 1E in the main text). Two of these are central (maroon beads in Fig. S1) and have a hard-core repulsion preventing the formation of two simultaneous 7S bonds within the same dimer. The other two pairs of patches (shades of blue and pink in Fig. S1) are such that blue patches attract blue patches and pick patches attract pink patches. Their positions and interaction-specificity guarantees that the stable bonded configuration is a bonded tetramer, with two protomers oriented in one direction and the remaining two in the opposite direction (as in Fig. 1B and E). This is inspired by *in vitro* observations of the structure of 7S tetramer bonds [2, 3].

Bonds result from the interaction of Eq. (S1), where the following parameters yield strong attraction and short-range repulsion: $\epsilon^{\text{7S}}_{\text{LJ}} = 15k_{\text{B}}T$, $r^{\text{7S}}_c = 3a$ and $a^{\text{7S}} = 0.2a$ for the blue and pink beads, and $\epsilon^{\text{7S}}_{\text{LJ}} = 2k_{\text{B}}T$, $r^{\text{7S}}_c = 3a$ and $a^{\text{7S}} = 0.4a$ for the maroon beads.

The NC1 domain is made up of one attractive bead surrounded by a ring of 6 repulsive beads of the same type as the chain, as in Fig. S1 and Fig. 1F. This ensures there is only space for one other NC1 end to bond. The central NC1 bead (maroon) is strongly attractive to beads of the same type, with short-range repulsion: $\epsilon^{\text{NC1}}_{\text{LJ}} = 20k_{\text{B}}T$, $r^{\text{NC1}}_c = 3a$ and $a^{\text{NC1}} = a$.

All beads of size $a$ have a mass of $m$, which we take as our simulation mass unit, and the patch beads of the 7S

| Parameter | Model value |
|---|---|
| $\kappa_{\mathrm{len}}$ | $50k_{\mathrm{B}}T/a^2$ |
| $r_0$ | $0.5a$ |
| $\kappa_{\mathrm{ang}}$ | 6 and $0.6k_{\mathrm{B}}T$ |
| $\beta_0$ | $180^\circ$ |
| $\epsilon_{\mathrm{LJ}}^{\mathrm{chain}}$ | $2k_{\mathrm{B}}T$ |
| $r_{\mathrm{c}}^{\mathrm{chain}}$ | $2^{1/6}a$ |
| $\epsilon_{\mathrm{LJ}}^{\mathrm{NC1}}$ | $20k_{\mathrm{B}}T$ |
| $a^{\mathrm{NC1}}$ | $a$ |
| $r_{\mathrm{c}}^{\mathrm{NC1}}$ | $3a$ |
| $\epsilon_{\mathrm{LJ}}^{\mathrm{7S}}$ | $15k_{\mathrm{B}}T$ |
| $a^{\mathrm{7S}}$ | $0.2a$ |
| $r_{\mathrm{c}}^{\mathrm{7S}}$ | $3a$ |
| $\Delta t$ | $0.002\mathcal{T}_0$ |
| $N_{\mathrm{bt}}$ | 1024 |
| $L_x, L_y, L_z$ | $64a$ |
| $m_{\mathrm{bead}}$ | $1m$ |
| $m_{\mathrm{patch}}$ | $0.1m$ |
| $m_{\mathrm{tot}}$ | $37.6m$ |
| $\mathcal{L}$ | $15.5a$ |

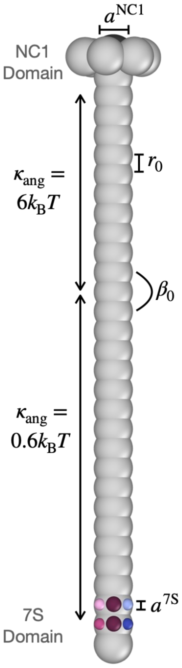


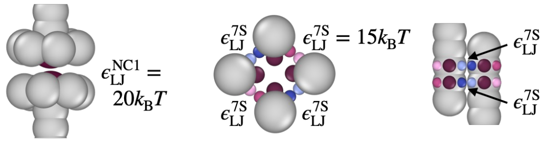


FIG. S1. **Molecular model.** Table with the model parameters and schematics. At the top right, a whole protomer. At the bottom, an NC1 bond (left), 7S bonds in a tetramer (center), and a 7S bond in a dimer (right).

domain have a mass of $0.1m$. All model parameters are summarized in Fig. S1.

**Simulation protocol** $N_{\mathrm{m}} = 1024$ protomers are simulated in a cubic box of length $L_{\mathrm{box}} = 64a$ with periodic boundary conditions. This is at the upper limit of what is tractable due to the large computational cost of this model. The simulations are performed in the canonical ensemble: a Velocity-Verlet integrator with timestep $\Delta t = 0.002\mathcal{T}_0$ is coupled to a Langevin thermostat [4] with a viscous friction coefficient $\gamma = m/\mathcal{T}_0$ for the chain beads and $0.1m/\mathcal{T}_0$ for the attractive patches. Here, $\mathcal{T}_0 = \sqrt{ma^2/(k_{\mathrm{B}}T)}$ is the simulation time unit. To simulate the assembly, stretch and relaxation of a network takes approximately 7 days.

After an equilibration over a time $6 \times 10^7\Delta t$, during which protomers form a connected network and network properties are no longer changing, strain is applied over a time $6 \times 10^6\Delta t$. The time over which the networks are strained is chosen to be short compared to the relaxation timescales, mimicking experiments using tissue stretchers [5]. The strain is applied by expanding the simulation box in the $z$-direction and simultaneously shrinking the simulation box in the $x$ and $y$ directions so that the total volume stays constant. See section II for a description of the strain protocol. After reaching the final strain, the network is left to re-equilibrate at this strain for $3.4 \times 10^7\Delta t$.

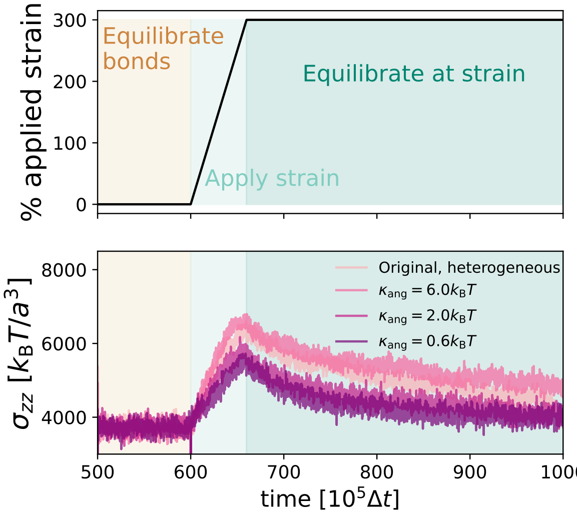


FIG. S2. **Effect of different bending coefficients in the molecular model.** There is evidence that collagen-IV has a heterogeneous persistence length [1], which informs our choice for a heterogeneous bending coefficient in the molecular model; a third of the chain beads close to the NC1 domain have a bending coefficient of $6.0k_{\mathrm{B}}T$ and the other two thirds have a bending coefficient of $0.6k_{\mathrm{B}}T$. This figure compares the original model to additional data of the stress over time when the bending coefficient is homogeneous along the chain and takes a value of $6.0k_{\mathrm{B}}T$, $2.0k_{\mathrm{B}}T$ and $0.6k_{\mathrm{B}}T$. The mechanics is not significantly affected by the different $\kappa_{\mathrm{ang}}$.

**Bond remodelling** When mutually attractive patches are within attraction range, a bond is formed. Likewise, bonds are broken via thermal fluctuations. Hence, the strength of the attraction determines the bond remodelling dynamics.

##### B. Bond topology model

**Model** In the bond topology model, a protomer is represented by a rigid rod connecting the 7S and NC1 ends. The rod has length $L_0 = 3a$ where $a$ is the simulation length unit. Each end forms harmonic bonds (NC1 or 7S) with the same end of other protomers. The bond energy is $E_{\mathrm{bt}} = \kappa(r-r_0)^2 - E_{\mathrm{bond}}$ where $r_0 = 0.5a$, $\kappa = 6k_{\mathrm{B}}Ta^{-2}$ and $E_{\mathrm{bond}}$ is varied between 3 and $20k_{\mathrm{B}}T$. In this model the length of a collagen-IV protomer within the network maps to the length of the rigid rod plus two half bonds, so that the length of a protomer is $L = L_0 + r_0 = 3.5a$. Since experimentally $L = 400$ nm [1], $a \approx 114$ nm in this model. We introduce a small penalty for the formation of acute angles between bonded protomers, implemented via a harmonic angular potential between protomers con-

| Parameter | Model value |
|---|---|
| $\kappa$ | $6.0k_\mathrm{B}T/a^2$ |
| $r_0$ | $0.5a$ |
| $\kappa_\mathrm{at}$ | $1.0k_\mathrm{B}T$ |
| $\beta_0$ | $180°$ |
| $\beta_0^{7\mathrm{S}}$ | $60°, 155°$ |
| $\Delta t$ | $0.002\mathcal{T}_0$ |
| $N_\mathrm{bt}$ | 3125 |
| $L_x, L_y, L_z$ | $12a$, $72a$, $72a$ |
| $m_\mathrm{tot}$ | $2m$ |
| $\mathcal{L}$ | $3.5a$ |

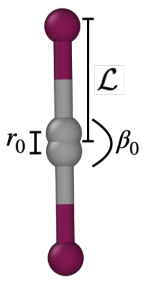


FIG. S3. **Bond topology model.** Table with the model parameters and a sketch of two protomers forming an NC1 bond (NC1 domain in gray, 7S domain in maroon).

nected via a bond: $E_\mathrm{at} = \kappa_\mathrm{at}(\beta - \beta_0)^2$ where $\beta$ is the angle between two bonded protomers. For bonded NC1 bonds $\beta_0 = 180°$, see Fig S3. Multiple angles are made when more than two 7S ends bond. These are the inner angle of the bond which we assign $\beta_0^{7\mathrm{S}} = 60°$, and the outer angle which connects to the rest of the protomer which we assign $\beta_0^{7\mathrm{S}} = 155°$. $\kappa_\mathrm{at} = 1k_\mathrm{B}T$ for all angle bonds. As detailed below, this prevents network collapse.

When two NC1 ends form a bond, they are prevented from forming additional bonds. Instead, the 7S end bonds with other 7S ends via a sequential reaction pathway (Fig. 2E): two 7S ends may bond with each other to form a dimer; then a third may bond with the dimer to form a trimer where all three 7S ends are connected by three bonds; finally, a fourth may bond the trimer, forming a tetramer with 6 bonds, where each protomer is connected to every other protomer. We limit ourselves to this sequential pathway as it has been previously suggested that two dimer bonds cannot come together to form a tetramer [6]. Other than NC1 and 7S bonds, there are no other interactions between protomers and, in particular, no steric repulsion is present. This makes the topology of the bonding pattern the characteristic of this model, hence its name. All beads have mass $m$. Model parameters are summarized in Fig. S3.

**Parameters choice** There are relatively few parameter choices defining the protomers in the bond topology model; here we briefly describe the rationale behind those choices. $\kappa$ is the bond constant dictating the longitudinal stiffness of the bonds. The exact value of $\kappa$ does not qualitatively affect the results but changes the phase space of the parameters in tandem with the strain applied. This can be seen in Fig. S5 where the stress and alignment relaxation is qualitatively the same for $\kappa = 6k_\mathrm{B}Ta^{-2}$, strain = 150% and $\kappa = 1k_\mathrm{B}Ta^{-2}$, strain = 260%. This is expected; the stress response is dictated by the initial elasticity of the network and then the likelihood of a bond breaking so that for a network with smaller $\kappa$ and therefore softer bonds, a larger strain is required for the same initial increase in stress and the same bond breaking dynamics.

The angular bond constant, $\kappa_\mathrm{at}$ is more constrained. There is no volume exclusion in the bond topology model, and so the angular bond constant must be large enough to prevent the networks from collapsing into small clusters (Fig. S4 shows the clustering taking place for $\kappa_\mathrm{at} = 0.1k_\mathrm{B}T$ and the absence of clustering for $\kappa_\mathrm{at} = 1k_\mathrm{B}T$). At the same time, $\kappa_\mathrm{at}$ should be small enough, such that the orientations of two bonded protomers are practically uncorrelated, as the persistence length of collagen-IV protomers is smaller than their length. We choose the value $\kappa_\mathrm{at} = 1k_\mathrm{B}T$, which avoids clustering while giving negligible alignment correlation along the network.

**Simulation protocol** $N_\mathrm{bt} = 3125$ protomers are placed in a simulation box the shape of a thin slab with dimensions $L_x = 12a$, $L_y = L_z = 72a$. We note that the mechanics of the system does not change significantly for larger systems, as shown in Fig. S6, where we increase the system size by a factor of 5. Periodic boundary conditions apply in the $y$ and $z$ dimensions, while Lennard-Jones walls delimit the $x$-boundaries. A Velocity-Verlet integrator with timestep $\Delta t = 0.002\mathcal{T}_0$ is coupled to a Langevin thermostat [4] with friction coefficient $\gamma = 10m/\mathcal{T}_0$, where $\mathcal{T}_0 = \sqrt{ma^2/(k_\mathrm{B}T)}$ is the simulation time unit. To simulate the assembly, stretch and relaxation of a network takes approximately 1 day.

The protomer network is allowed to equilibrate in the cubic box until network properties are no longer changing which takes between $3 - 20 \times 10^6\Delta t$ depending on the bond remodelling parameters. Then a linear strain is applied over a time $5 \times 10^4\Delta t$ and finally the network is left to re-equilibrate at this strain for a time which varies between $2 - 10 \times 10^6\Delta t$ depending on the bond remodelling parameters. Strain is applied by expanding the simulation box in the $z$-direction and keeping the $x$ and $y$ dimensions constant, see section II for further details.

**Bond remodelling** To assess the effect of non-equilibrium processes on the network mechanics, we compare equilibrium and out-of-equilibrium bond remodeling mechanism. In sections Results A and B, the bonds are remodelled to locally satisfy detailed balance [7]. Bonds are proposed to be made/broken at a rate $k_\mathrm{bond} = 0.001(\Delta t)^{-1}$ and are made/broken such that Eq. (1) is satisfied with $P_\mathrm{make}^\mathrm{EQ} = 0.1$, where $\Delta E$ is the difference in energy of the bond, $E = \kappa(r - r_0)^2 - E_\mathrm{bond}$. The energy $E_\mathrm{bond}$ is varied for different bond remodelling dynamics, so that protomers with large $E_\mathrm{bond}$ will form networks whose bonds rarely break, and protomers with small $E_\mathrm{bond}$ form networks whose bonds break often.

In section Results C, the bond remodelling is instead not constrained by detailed balance and therefore out of equilibrium, mimicking the action of enzymes. We choose a protocol where bonds are made/broken according to a

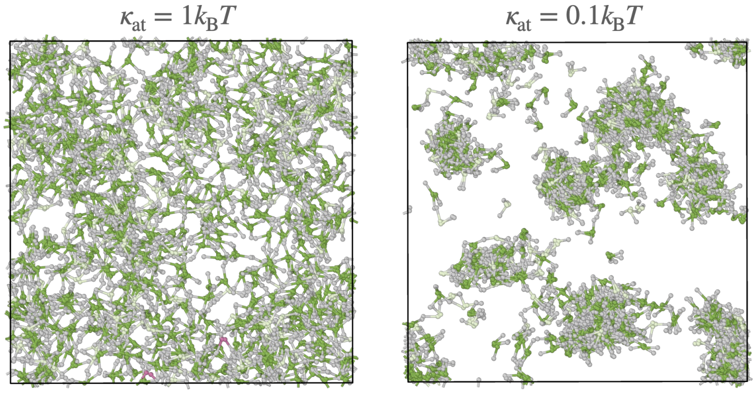


FIG. S4. **Effect of varying $\kappa_{\mathrm{at}}$.** In the bond topology model, the parameter $\kappa_{\mathrm{at}}$ controls both i) the correlation of the alignment of consecutive bonded protomers, giving rise to an effective non-zero persistence length within the network, and ii) how clustered the networks are. A value of $\kappa_{\mathrm{at}} = 1k_{\mathrm{B}}T$ is chosen such that box-spanning networks form yet protomers are flexibly bonded. These simulations were performed in the equilibrium bond remodelling regime with $E_{\mathrm{bond}} = 7k_{\mathrm{B}}T$, see section B. Protomer color corresponds to Fig. 2E in the main.

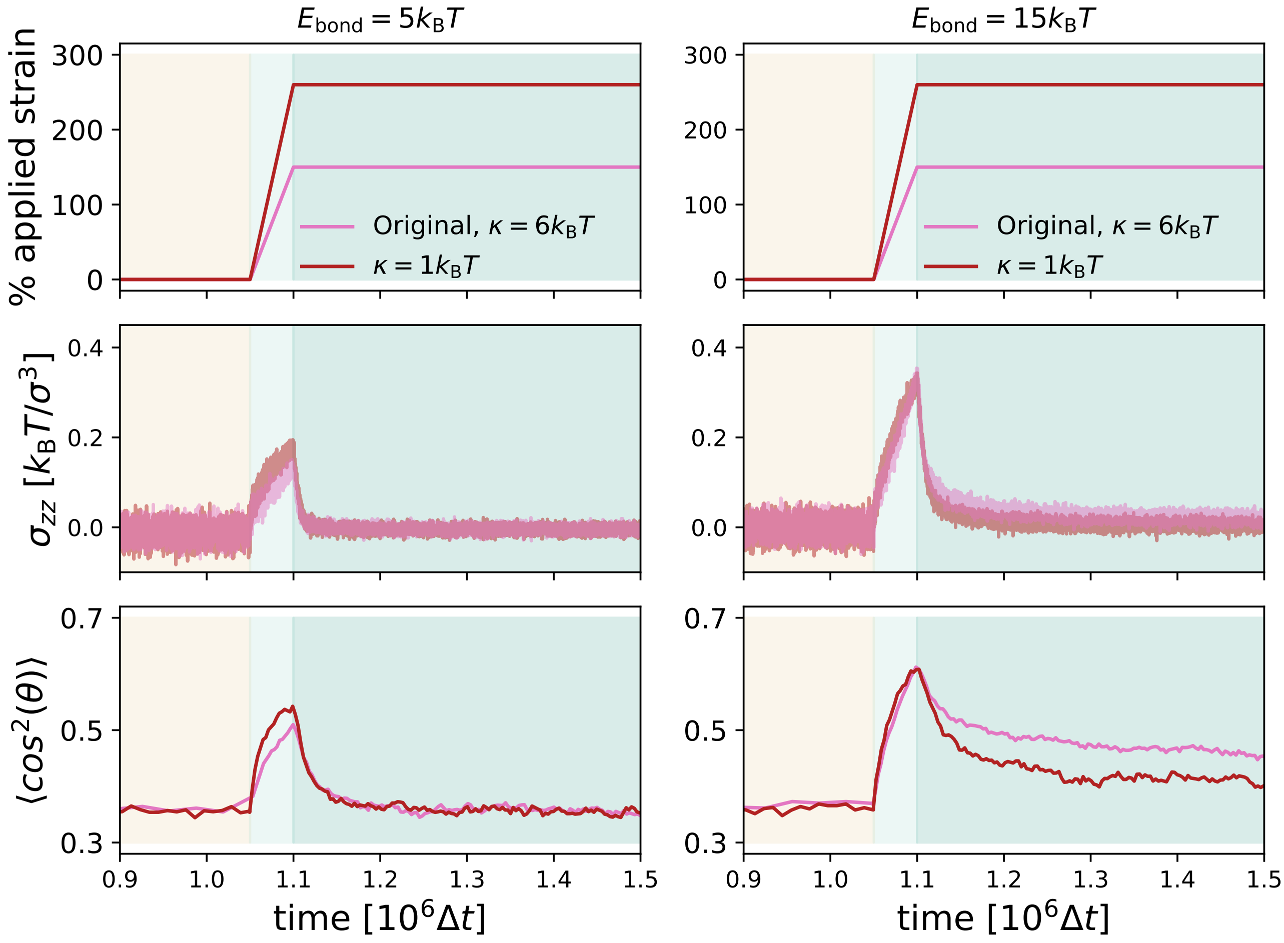


FIG. S5. **Effect of bond elastic constant.** The exact value of the bond constant, $\kappa$, which determines the energy to stretch a bond longitudinally in the bond topology model, does not qualitatively influence the mechanics of the networks. For two different values of $\kappa$ ($\kappa = 6.0k_{\mathrm{B}}T/a^2$, pink and $\kappa = 1.0k_{\mathrm{B}}T/a^2$, red) the stress and alignment is measured. The mechanics are equivalent for shifted values of the applied strain, where a smaller $\kappa$ requires a larger strain to exhibit the same mechanics as the larger $\kappa$ networks.

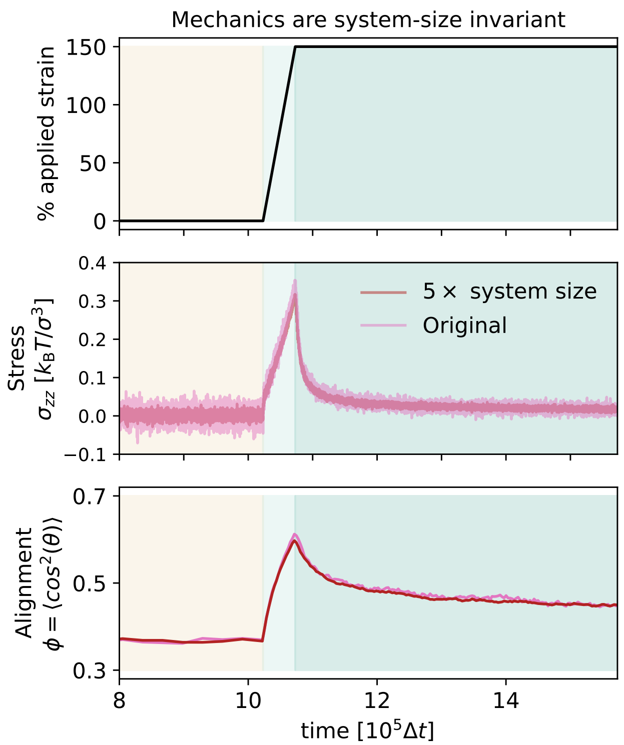


FIG. S6. **Effect of change in system size.** In the bond topology model, the number of protomers is increased by $5\times$, keeping the density and box size in the x-direction, $L_{\mathrm{x}}$, constant. The mechanics is unaffected by system size. These simulations are in the equilibrium bond remodelling regime with $E_{\mathrm{bond}} = 15k_{\mathrm{B}}T$.

rate $C^{\mathrm{enzy}}_{\mathrm{break/make}}$ and an activation distance $D^{\mathrm{enzy}}_{\mathrm{break/make}}$: bonds are made with probability $C^{\mathrm{enzy}}_{\mathrm{make}}$ if two protomer ends are closer than the activation distance $D^{\mathrm{enzy}}_{\mathrm{make}}$ and broken with a probability $C^{\mathrm{enzy}}_{\mathrm{break}}$ if they are further away than the activation distance, $D^{\mathrm{enzy}}_{\mathrm{break}}$. In Fig. S10 we explore the relaxation of networks which follow the enzymatic bond remodelling protocol for $D^{\mathrm{enzy}}_{\mathrm{break}} = 0$, a bond of any length can break with probability $C^{\mathrm{enzy}}_{\mathrm{break}}$. These networks have very dynamic bond remodelling and are therefore susceptible to forming clusters, as in Fig. S4. For this reason, the data in Fig. S10 are for networks with $\kappa_{\mathrm{at}} = 4k_{\mathrm{B}}T$.

#### II. STRAIN PROTOCOL

A strain is applied to fully connected networks for both models in sections Results B and C of the main paper. This strain is implemented using the LAMMPS command `fix deform` [8, 9]. In the bond topology model, this command is used to apply a strain in the $z$-direction. The keyword `remap x` is specified so that all bead coordinates are remapped to new coordinates in the strain axis as the simulation box is deformed. The strain linearly increases over $10^4\Delta t$ such that the final strain is 150%, unless otherwise specified (see Fig. 3B). A similar protocol is used for the molecular model, with the only difference being the use of the keywords `y volume x volume` so that the box size decreases in the $x$ and $y$ directions as it is increased in $z$, to preserve total volume. In the molecular model, the strain is applied linearly over $60 \times 10^5\Delta t$ to a final strain of 300% (see Fig. 3A). Fig. S7 shows snapshots of the networks before and after strain is applied. The velocities of all beads are unchanged during the deformation.

#### III. CRITICAL STRAIN AND YOUNG MODULUS

We investigated the behaviour of our networks for small and slowly applied strains, within the bond topology model. This allowed us to extract the Young modulus and to estimate the critical strain, defined as the strain at which a floppy network rigidifies and starts stiffening [10–12]. To do so, we performed simulations without bond remodelling and at a strain rate 10 times smaller than in the original stretch protocol. Fig. S8 shows the resulting stress-strain curve.

From the linear regime at small strain we extracted an elastic modulus of $\approx 0.06k_{\mathrm{B}}Ta^{-3}$ in simulation units, corresponding to $\approx 0.03$ Pa at physiological temperature (black dotted line). The estimated critical strain can be seen when the stress-strain slope increases at $\Delta L^*/L_0 \approx 1.1$.

The Young modulus of our model networks is significantly lower than values measured for reconstituted basement membrane matrix, which are in the range of $50 - 500$ Pa [13]. However, the basement membrane contains a plethora of other molecules like laminin, perlecan and nidogen [14]. We suspect that at low strain, the mechanics of the basement membrane is likely dictated by the collagen-IV network connectivity, but also by other ingredients such as cross-linkers and the laminin network [15]. Additionally, the reported mechanical measurements on reconstituted basement membrane matrix might have been obtained from bulk samples, which would differ significantly from the properties of a thin sheet such as the one considered in this work. Finally, in the real system, the collagen-IV matrix might be prestressed during development, but since the amount of pre-stress is not known, this small-strain measurement on a network without pre-stress might not be physiologically relevant. These factors might explain the large discrepancy between our measured value and those reported in the literature.

#### IV. PROTOMER DIFFUSION TIMESCALE

In the simulations, the action of the solvent is modelled implicitly by two forces acting on each bead: one

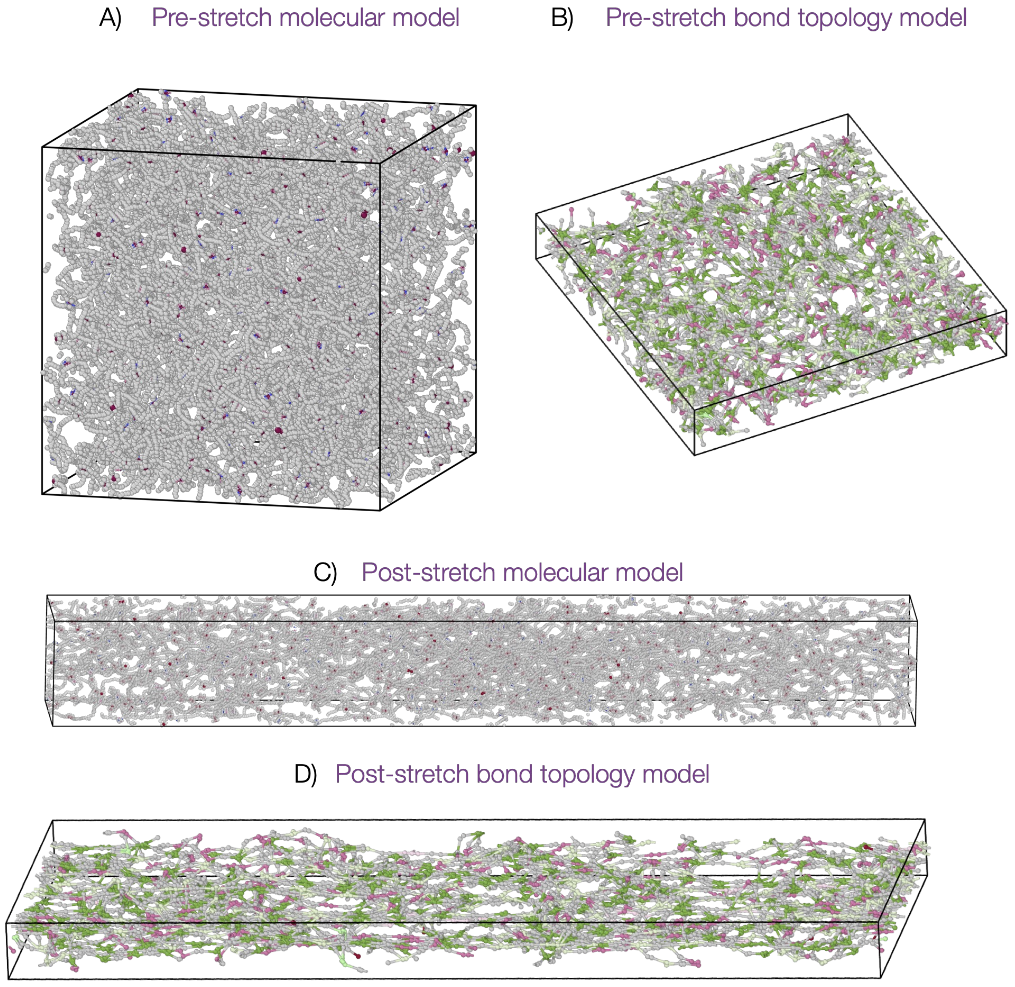


FIG. S7. **Network snapshots.** (**A**) and (**B**) snapshots of the molecular model and bond topology model, respectively, before stretching. (**C**) and (**D**) snapshots of the molecular model and bond topology model, respectively, immediately after stretching. The strain is 300% for the molecular model and 150% for the bond topology model. The colors of the molecules correspond to those in Fig. 1D and Fig. 2E of the main for the molecular model and bond topology model, respectively

represents the viscous friction (with friction coefficient $\gamma$) and the other the random collisions with the solvent molecules. In this model, a molecule of $n$ beads will experience a total friction $n\gamma$, as in the Rouse model for polymers [16]. Thus, for a protomer of $n$ beads, we have a diffusion coefficient

$$D = \frac{k_B T}{n\gamma}. \tag{S2}$$

From this relation, we can estimate the time a protomer takes to diffuse over its own size. In the bond topology model, a protomer is represented by two beads connected by a rigid linker of length $L = 3a$, with $a$ the bead diameter. Thus, $n = 2$ and the molecular size is $R = L = 3a$. The diffusion time scale for the bond topology model is thus

$$\tau_{\mathrm{BT}} = \frac{R^2}{D} = \frac{9a^2\gamma}{2k_B T} = 45\mathcal{T}_0 = 4.5 \times 10^3 \Delta t, \tag{S3}$$

since $\gamma = 10m/\mathcal{T}_0$ and $\Delta t = 2 \times 10^{-3}\mathcal{T}_0$ for this model.

For the molecular model, we can take as a measure of the molecular size the RMS end-to-end distance $\sqrt{\langle R^2 \rangle}$ of the protomer. We measured this quantity directly from the configurations, finding $\sqrt{\langle R^2 \rangle} = 8.3a$. Since the total number of beads in a protomer including the NC1 domain is $n = 37$, the diffusion time can be evaluated as

$$\tau_{\mathrm{molecular}} = \frac{\langle R^2 \rangle}{D} = \frac{\langle R^2 \rangle n\gamma}{k_B T} = 2553\mathcal{T}_0 = 1.3 \times 10^6 \Delta t, \tag{S4}$$

since $\gamma = m/\mathcal{T}_0$ and $\Delta t = 2 \times 10^{-3}\mathcal{T}_0$ for this model. In computing the diffusion coefficient of the protomer,

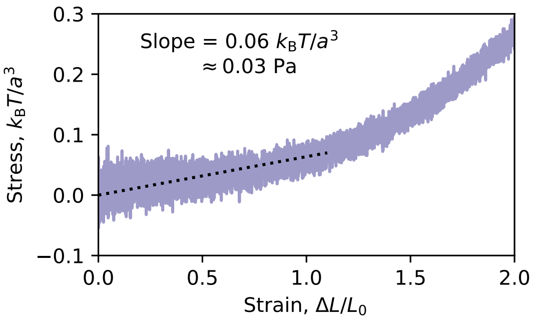


FIG. S8. **Estimating critical strain and Young modulus.** Stress-strain curve for the bond topology model at 10 times slower strain rate than in the original stretch protocol. The dashed black line is a linear fit of the strain-stress before strain stiffening, giving an estimate for the Young modulus in the linear elastic regime of 0.03Pa. The critical strain, the onset of strain stiffening, is roughly 1.1.

we have neglected the contribution from the attractive patches, which experience a friction 1/10 of the one experienced by the other beads.

By imposing that times $\tau_{\text{molecular}}$ and $\tau_{\text{BT}}$ represent the same physical time, we obtain that the time units $\mathcal{T}_0$ of the molecular and bond topology models should be in a ratio $2553/45 = 57$. Analogously, the simulation time step $\Delta t$ in the bond topology model should represent a time 290 times larger than in the molecular model. This tentative mapping might not hold exactly, especially at small times and during percolation, both because of the $k_{\text{bond}}$-limited remodelling rate in the bond topology model and because binding in the molecular model might not require diffusion over the whole protomer size [17]. Note also that the results we present are obtained at slightly higher density for the molecular than for the bond topology model (2.3 vs. 1.3 protomers/$R^3$). Regardless, this computation shows that the more coarse grained model is potentially two orders of magnitude faster.

#### V. PERSISTENCE LENGTH OF THE CHAIN (MOLECULAR MODEL)

In the molecular model, a collagen-IV protomer is represented as a chain of 30 beads connected to 7 additional beads that represent the NC1 domain.

We set $\kappa_{\text{ang}} = 6k_BT$ for the 11 triplets close to the NC1 domain, and $\kappa_{\text{ang}} = 0.6k_BT$ for the 17 remaining triplets.

To measure the average persistence length of the protomer, we consider the 30 beads of the chain plus the central bead of the NC1 domain, connected to these. The persistence length can be obtained from average cosine of the triplet angle $\beta$ [18]: $L_p = -r_0/\log[\langle\cos(\beta)\rangle]$ ($r_0$=bond length). We find $L_p = 1.80a$ if the whole chain is considered, and $L_p = 1.15a$ if the last, stiffer, 11 triplets are excluded.

#### VI. SUPPLEMENTARY VIDEOS

**Video 1:** This video depicts the self-assembly of networks in the bond topology model, from the top-down view of the $yz$ plane. The colours of the protomers change when they bond into dimers, trimers and tetramers according to the colour scheme in Fig. 2E. The bond strength here is $E_{\text{bond}} = 15k_BT$ and the equilibrium remodelling protocol is employed.

**Video 2:** This video depicts the self-assembly in the Molecular model. The protomers are coloured as in Fig. 1D. The bond strengths are $E_{\text{LJ}}^{NC1} = 20k_BT$ and $E_{\text{LJ}}^{NC1} = 15k_BT$.

**Video 3:** This video depicts the stretching and subsequent network relaxation of networks in the Bond Topology model from the top-down view of the $yz$ plane. The parameters are the same as Video 1. The strain applied is 150%.

**Video 4** This video depicts the stretching and subsequent network relaxation of networks in the Molecular model. The parameters are the same as Video 2. The strain applied is 300%.

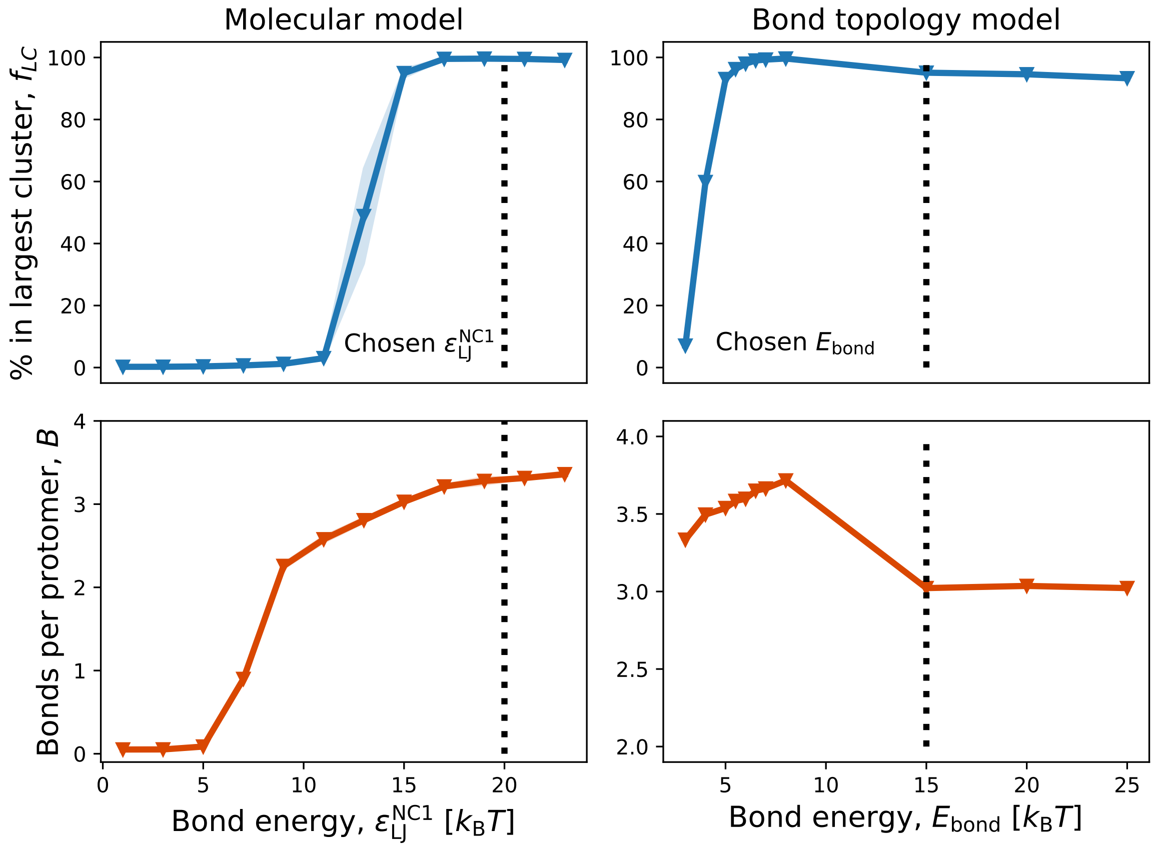


FIG. S9. **Network properties as a function of protomer-protomer bond energy.** (**A**) and (**B**) the fraction of protomers in the largest connected network as a function of protomer-protomer bond strength showing that the networks percolate at sufficient bond energy. The bond strength in the molecular model as defined by $\epsilon_{LJ}^{NC1}$, where $\epsilon_{LJ}^{7S}$ is changed proportionally according to the table in Fig. S1. The bond strength in the bond topology model as defined by $E_{bond}$. The bond energies used in the subsequent work in Figs. 2 and 3 are $20k_BT$ for the molecular model and $15k_BT$ in the bond topology model, as indicated. (**C**) and (**D**) the bonds per protomer saturate to $<4$ for both the molecular model and the bond topology model as bond energy is increased. The bond topology model shows an additional regime at high bond energy whereby the average bonds per protomer decreases. This is due to the networks becoming kinetically trapped in a non-optimal bonding configuration.


[1] A. Al-Shaer, A. Lyons, Y. Ishikawa, B. G. Hudson, S. P. Boudko, and N. R. Forde, Biophysical Journal **120**, 4013 (2021).
[2] B. Siebold, R.-q. Qian, R. W. Glanville, H. Hofmann, R. Deutzmann, and K. Kähn, European Journal of Biochemistry **168**, 569 (1987).
[3] K. Kühn, H. Wiedemann, R. Timpl, J. Risteli, H. Dieringer, T. Voss, and R. W. Glanville, FEBS Letters **125**, 123 (1981).
[4] T. Schneider and E. Stoll, Physical Review B **17**, 1302 (1978).
[5] M. Duda, N. J. Kirkland, N. Khalilgharibi, M. Tozluoglu, A. C. Yuen, N. Carpi, A. Bove, M. Piel, G. Charras, B. Baum, and Y. Mao, Developmental Cell **48**, 245 (2019).
[6] P. D. Yurchenco and H. Furthmayr, Annals of the New York Academy of Sciences **460**, 530 (1985).
[7] M. E. Tuckerman, *Statistical mechanics: theory and molecular simulation* (Oxford university press, 2023).
[8] A. P. Thompson, H. M. Aktulga, R. Berger, D. S. Bolintineanu, W. M. Brown, P. S. Crozier, P. J. in 't Veld, A. Kohlmeyer, S. G. Moore, T. D. Nguyen, R. Shan, M. J. Stevens, J. Tranchida, C. Trott, and S. J. Plimpton, Comp. Phys. Comm. **271**, 108171 (2022).
[9] Lammps: fix deform, `https://docs.lammps.org/fix_deform.html`, accessed: 2024-07-16.
[10] A. J. Licup, S. Münster, A. Sharma, M. Sheinman, L. M. Jawerth, B. Fabry, D. A. Weitz, and F. C. MacKintosh, Proceedings of the National Academy of Sciences **112**, 9573 (2015).
[11] A. Sharma, A. J. Licup, K. A. Jansen, R. Rens, M. Sheinman, G. H. Koenderink, and F. C. MacKintosh, Nature Physics **12**, 584 (2016).
[12] C.-T. Lee and M. Merkel, Soft Matter **18**, 5410 (2022).

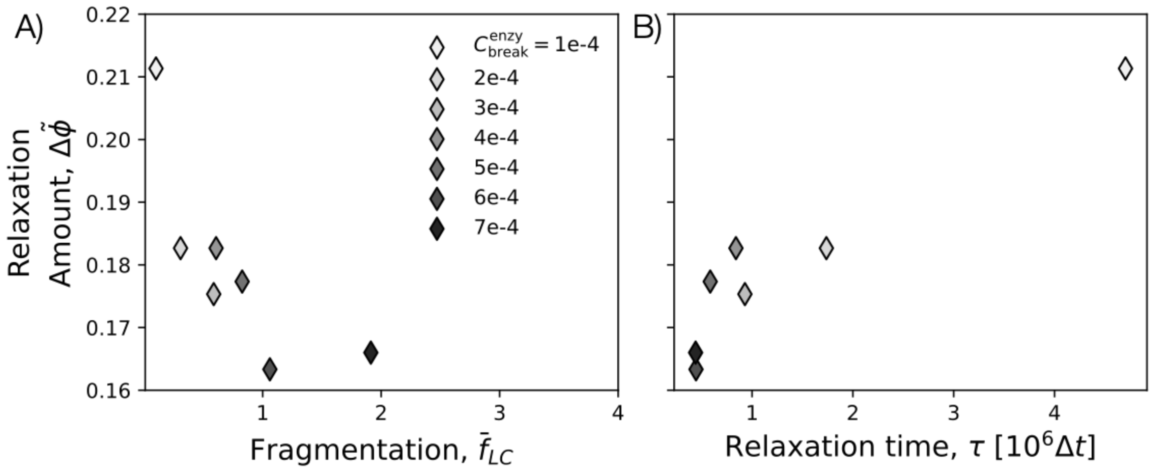


FIG. S10. **Non mechano-sensitive enzymatic bond remodelling.** (**A**) and (**B**) show the relaxation properties of networks whose bonds remodel in a non mechano-sensitive way. A bond of any length can break, with the probability shown in the legend. (**A**) the relaxation amount scaled by strain, $\Delta\tilde{\phi}$, takes a variety of values while the fragmentation, $\bar{f}_{\rm LC}$ stays low. (**B**) the relaxation timescale, $\tau$, also takes a variety of values. For these networks, $D_{\rm make} = 0.65a$ and $C_{\rm make} = 1.0$, see SI section IB for further details on the non mechano-sensitive simulation setup.

[13] O. Chaudhuri, J. Cooper-White, P. A. Janmey, D. J. Mooney, and V. B. Shenoy, Nature **584**, 535 (2020).
[14] R. Jayadev and D. R. Sherwood, Current Biology **27**, R207 (2017).
[15] N. Khalilgharibi and Y. Mao, Open Biology **11**, 200360 (2021).
[16] M. Rubinstein and R. H. Colby, *Polymer physics* (Oxford university press, 2003).
[17] P. G. de Gennes, The Journal of Chemical Physics **76**, 3316 (1982).
[18] H.-P. Hsu and K. Kremer, The Journal of Chemical Physics **144**, 154907 (2016).
